# Atypical neuromagnetic resting activity associated with thalamic volume and cognitive outcome in very preterm children

**DOI:** 10.1101/729038

**Authors:** Adonay S. Nunes, Nataliia Kozhemiako, Evan Hutcheon, Cecil Chau, Urs Ribary, Ruth E Grunau, Sam M Doesburg

## Abstract

Children born very preterm, even in the absence of overt brain injury or major impairment, are at risk of cognitive difficulties. This risk is associated with disruption of ongoing critical periods involving development of the thalamocortical system while in the neonatal intensive care unit. The thalamus is an important structure that not only relays sensory information but acts as a hub integrating cortical activity, and through this integration, it regulates cortical power at different frequency bands. In this study, we investigate the association between atypical power at rest in children born very preterm at school age, neurocognitive function and structural alterations related to the thalamus. Our results indicate that children born extremely preterm have higher power at low frequencies and lower power at high frequencies, compared to controls born full-term. A similar pattern of spectral power was found to be associated with poorer neurocognitive outcomes. This pattern of spectral power was also associated with normalized T1 intensity and the volume of the thalamus. Overall, this study provides evidence of the relation between structural alterations related to very preterm birth, atypical oscillatory power at rest and neurocognitive difficulties at school-age children born very preterm.

## Introduction

Worldwide, the incidence of preterm birth with a very low gestational age (VLGA, born ≤ 32 weeks of GA) is 1 in 100, and extremely low gestational age (ELGA, born ≤ 32 weeks of GA) 1 in 200, with a mortality rate of 5-10% and +10%, respectively [Torchin et al., 2015]. Even in the absence of overt brain lesions, very preterm infants are at higher risk of cognitive, behavioral and motor problems [Aarnoudse-Moens et al., 2009; Anderson, 2014; Moore et al., 2012] compared to healthy full-term children. Risk factors in very preterm infants (ELGA & VLGA) include post-natal infection [Ranger et al., 2015; Zwicker et al., 2016], as well as exposure to numerous daily invasive procedures such as blood draws and line insertions, during weeks to months in the neonatal intensive care unit (NICU) [Roofthooft et al., 2014; Vinall and Grunau, 2014]. The disruption of normal intrauterine maturation by preterm birth, neonatal clinical factors, and pain-related stress alters ongoing brain development during critical periods involving prolific axonal growth, dendritic sprouting, and synapse formation [Mrzljak et al., 1992], that leads to abnormal brain morphology and activity [Kapellou et al., 2006; Schneider et al., 2018; Smith et al., 2011]. Thus, to ensure proper extrauterine environment and adequate preterm interventions, it is of vital importance to understand how structural and functional preterm alterations are associated with cognitive and behavioral outcomes in children born very preterm.

During the neonatal period in very preterm infants born 24-32 weeks of gestation, thalamic afferents that are synapsing in the subplate, a transient structure that generates endogenous activity and critical to the formation of long term thalamocortical circuitry [Kostovic and Judas, 2002], gradually synapse onto cortical neurons and become sensory driven. Disruption in this developmental process might cause neural apoptosis [Anand et al., 2007; Dührsen et al., 2013] and altered development of thalamocortical axons [Dean et al., 2013; Molnar and Rutherford, 2013]. Moreover, very preterm infants present abnormal myelinization and fractional anisotropy in gray and white matter areas [Dubois et al., 2008; Eaton-Rosen et al., 2015], decreased cortical and subcortical gray matter volume [Dubois et al., 2008; Eaton-Rosen et al., 2015], decreased cortical and subcortical gray matter volume [Boardman et al., 2006; Chau et al., 2019], thalamocortical alterations [Ball et al., 2012; Cai et al., 2017] and pain-induced volume reduction of white matter and subcortical gray matter [Brummelte et al., 2012; Duerden et al., 2018]. These structural alterations persist into school age [Hohmeister et al., 2010; Lax et al., 2013] and adulthood [Menegaux et al., 2017; Nosarti et al., 2014] and are associated with lower IQ [Breeman et al., 2017; Nosarti et al., 2014].

The thalamus acts as a sensory relay by directing peripherical information to the cortex and as an integrative hub for cortical representations by mediating cortico-cortical communication [Hwang et al., 2017; Jones, 1998; Malekmohammadi et al., 2015; Sherman and Guillery, 2013]. It has been associated with numerous cognitive functions [Jones, 2012; Sherman, 2016; Theyel et al., 2010] and affected in many disorders [Giraldo-Chica et al., 2018; Llinás et al., 1999; Llinás et al., 2005; Ribary et al., 2019; Tona et al., 2014]. Functionally, the thalamus regulates cortical power at different frequencies mediating alpha and theta-gamma switching [Ribary et al., 2017]. Neurological conditions are often associated with the thalamus slowing alpha oscillatory activity to theta [Lianyang Li et al., 2015; Ribary et al., 2019; Tarapore et al., 2013], and in very preterm children at school age decreased alpha power and increased gamma have been reported [Cepeda et al., 2007; Doesburg et al., 2010; Doesburg et al., 2011], as well as atypical neurophysiological connectivity [Doesburg et al., 2013; Kozhemiako et al., 2019; Moiseev et al., 2015b].

Based on previous literature reporting associations between the thalamus and cortical power [Edgar et al., 2015; Lindgren et al., 1999; Ribary et al., 2017], in the present study, we hypothesized that reductions in thalamic volume present in children born very preterm will be associated with atypical cortical power measured using MEG. Given that power captured by MEG is mostly generated by pyramidal neurons in the cortex [Murakami and Okada, 2006], to assess the association between thalamic volume and power at different frequencies, cortical gray matter volume was included as a variable of interest, and both were corrected for total intracranial volume. In addition, previous studies have suggested that T1 intensities reflect myelinization [Koenig, 1991; Stüber et al., 2014], gray matter intensity has been used as a biomarker for neurological diseases and aging [Kong et al., 2015; Norbom et al., 2018; Salat et al., 2009], and it has been reported myelin alterations in the thalamus and cortical gray matter in very preterm infants [Counsell et al., 2002; Counsell et al., 2014; Eaton-Rosen et al., 2015]. Thus, a separate association analysis was conducted to test if mean thalamic, cortical gray matter and cortical white matter T1 intensities are associated with frequency-specific neurophysiological power at rest.

The aim of the present study is to investigate the relationship between structural alterations in very preterm children at school age and atypical oscillatory power at canonical frequency bands, and their association with neurodevelopment at school age. To this end, using a multivariate statistical analysis we estimated differences in MEG relative power at canonical frequency bands between ELGA, VLGA and Term born children, the association between power and cognitive, visual-motor and behavior problems at school age, and the association between power and thalamic/cortical volumes or intensities measured using MRI.

## Methods

### Participants

A total of 108 children participated in a resting state MEG recording, from which 9 participants were discarded due to major brain injury (periventricular leukomalacia [PVL] or grade III-IV intraventricular hemorrhage [IVH] on neonatal ultrasound), autism spectrum disorder [ASD], and/or or excessive motion. The final sample size was 99 children with a mean age of 7.8 years: 23 were born ELGA (24 to 28 wks, age 7.7 ± 0.39, 10 girls), 36 VLGA (29–32 wks GA, age 7.7 ± 0.39, 24 girls), and 39 healthy full-term (40 wks GA, age 7.9 ± 1.02, 24 girls). Of these 99 children, only 62 underwent MRI, and after discarding scans with poor quality, 51 participants remained, consisting of 13 ELGA, 24 VLGA and 13 full-term children. All children were recruited as part of a prospective longitudinal study on the effects of neonatal pain-related stress on neurodevelopment of children born very preterm [Grunau et al., 2007; Grunau et al., 2009]. Participant characteristics are presented in Table 1.

**Table 1.**
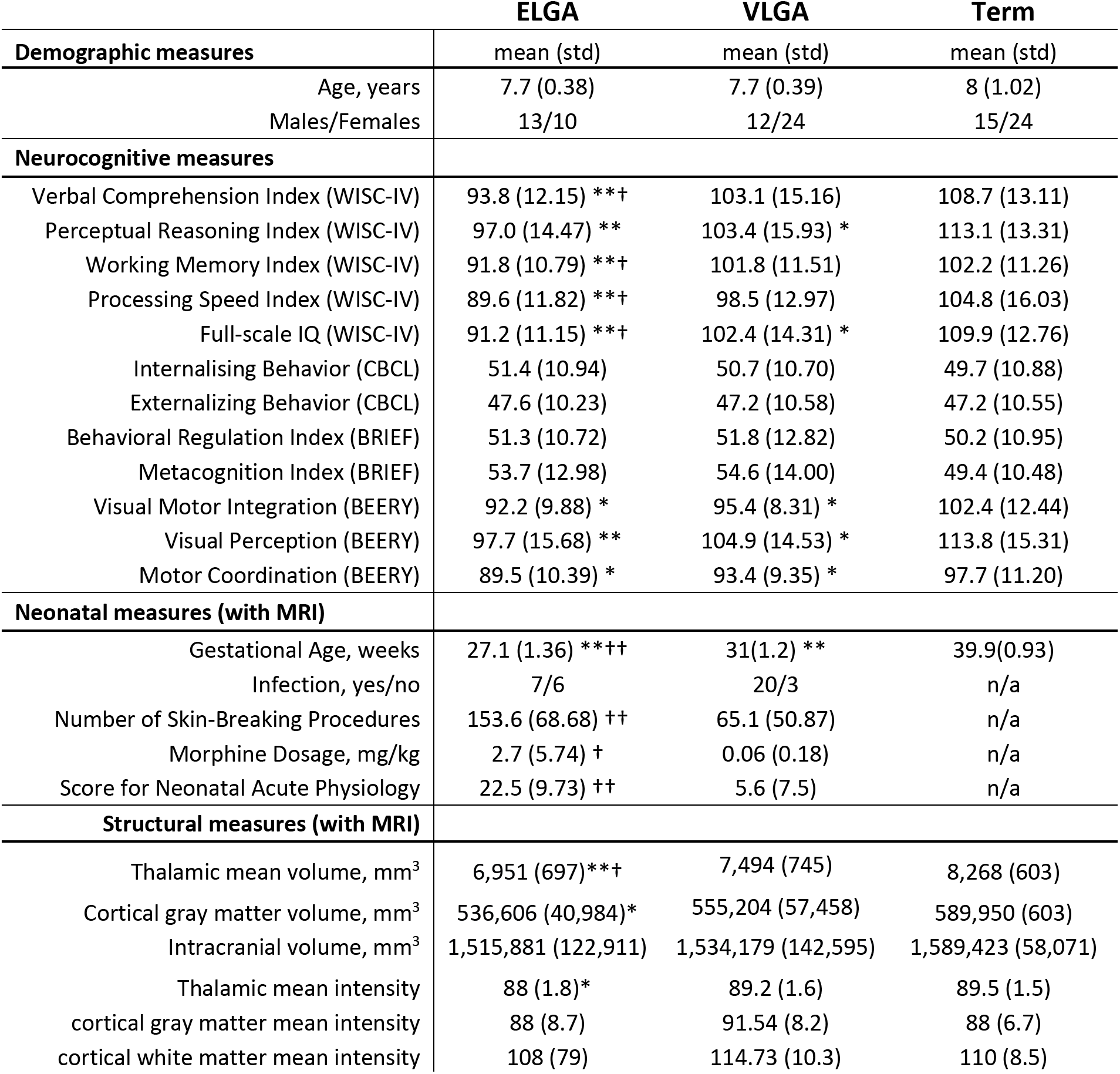
Study cohort characteristics. Significant differences between ELGA or VLGA and full-term *p ≤.05,**p ≤.001. Significant differences between ELGA and VLGA †p ≤.05, †† p ≤.001.

### MEG recording

Eyes open resting state MEG data were recorded for a total of two minutes using a CTF 151-Channel MEG system (CTF systems; Coquitlam, Canada) in a magnetically shielded room with a sampling rate of 1200 Hz. Continuous head localization was recorded by energizing three fiducial coils placed in the nasion and preauriculas. The head shape of the participants was recorded using a Polhemus Fastrak digitizer.

### MRI recording

MRI scans were performed on a Siemens 1.5 Tesla Avanto (Berlin, Germany) using a 12 channel head coil. Each scan consisted of a 3D T1-weighted sequence (TR:18; TE:9.2; FOV:256; Thickness:1mm; Gap:0; Matrix:256×256).

### Neonatal data

Neonatal data were collected from a daily chart review by an experienced research nurse during the neonatal period of the ELGA and VLGA participants as described previously [Grunau et al., 2009]. In this study, we focused on gestational age (GA), if infection was positive, illness severity on day 1 (SNAP; [Richardson et al., 2001]), equivalent log-transformed cumulative morphine dose, and log-transformed number of skin-breaking procedures (pain), from birth to term-equivalent age.

### Neurocognitive assessment

On the day of the MEG scanning, child IQ was assessed with the Wechsler Intelligence Scale for Children (WISC-IV; Wechsler, 2003), yielding standardized scores for verbal (VIQ), perceptual reasoning (IQ-PR), working (IQ-WM), processing speed (IQ-PSI) and full-scale (IQ-f). Visual-motor capabilities were assessed with Beery–Buktenica Developmental Test of Visual-Motor Integration, 5th Edn. [Beery et al., 2004], comprising the subscales visual-motor integration (BEERY-VMI), visual perception (BEERY-VP), and motor coordination (BEERY-MC). Behavior was assessed with the Child Behavior Checklist (CBCL; Achenbach and Rescorla 2001) questionnaire completed by a parent, measuring internalizing behavior (CBCL-INT and externalizing behavior (CBCL-EXT), and executive functions with the Behavior Rating Inventory of Executive Function (BRIEF; [Gioia et al., 2000] assessing behavioral regulation (BRIEF-BRI) and metacognition index (BRIEF-MCI).

### MEG analysis

MEG sensor data were notch filtered at 60 Hz to remove line noise. Segments exceeding 5 mm displacement at any direction from the median head position were discarded. After visual inspection, segments with muscle artifacts were also discarded. Independent Component Analysis (ICA) was used to identify and reject components capturing eye and heart activity. The remaining artifact-free data was segmented into epochs of four seconds and band-pass filtered at canonical frequency bands (delta:1-4 Hz, theta:4-8 Hz, alpha:8-12 Hz, beta:12-25 Hz, gamma:25-55 Hz). For source reconstruction, a forward model was computed using a single-shell head model [Nolte, 2003]. For subjects without an MRI, the best match from a pool of child MRIs was used instead [Gohel et al., 2017]. Source space activity was reconstructed on an 8 mm spaced grid using an LCMV beamformer [Van Veen et al., 1997] which creates spatial filter weights that maximize activity from the target location while filtering out activity from elsewhere. Beamformer weights were estimated at each frequency band to optimize the spatial filtering [Moiseev et al., 2015a]. For each reconstructed source, relative power was obtained by dividing the absolute power within a frequency band by the sum of the absolute power in all the frequencies. Relative power is used as it is corrected for depth bias and it has been shown to be more stable across subjects [Suppiej et al., 2017]. The relative power within a frequency represents its contribution to all the frequency spectrum of interest. Herein, power and relative power is used interchangeably.

### Morphometric measures

3D T1 MRIs were automatically processed using FreeSurfer [Fischl, 2012], in brief, steps included skull-stripping, Talairach transformation, intensity normalization, subcortical segmentation, tessellation of the gray matter/white matter boundary, topology correction and surface deformation to detect gray matter/white matter and gray matter/cerebrospinal fluid boundaries [Fischl et al., 1999]. For this study, the volumetric measures of interest were total cranial volume, cortical gray matter volume and thalamic volume, and the intensity measures of interest were the mean normalized intensity of the thalamus, cortical gray matter and cortical white matter (sampled normal at 2mm under the cortical surface). Several studies report gray matter intensities contrasted with white matter intensities [Jefferson et al., 2015; Kong et al., 2015; Salat et al., 2009], while preserving the TI, to account for white matter intensity, it was included as a separate variable in the correlation analysis with power.

### Statistical analyses

Partial Least Squares (PLS) and linear regression were used for statistical analysis. PLS is a multivariate technique used in neuroimaging [Krishnan et al., 2011; McIntosh and Lobaugh, 2004] based on singular value decomposition. It decomposes the data into latent variables (LV) composed of a left singular vector (left-SV), a singular value and a right-SV. Permutation is used to assess statistical significance by resampling without replacement the subjects’ group assignment and a p-value is obtained by counting the number of permutations where the permutation singular value exceeded the original singular value. PLS yields a single p-value for each LV, and accordingly does not require correction for multiple comparisons. Bootstrapping is used to measure the reliability of the features by resampling with replacement subjects within a group and taking the standard error (SE) of the bootstrapped left-SVs, then the original left-SV is divided by the bootstrap SE, and a bootstrap ratio is obtained. This ratio is presented as z-scores. The right-SV is termed the contrast, as it reflects the data-driven contribution of the groups or variables to the right-SV.

In this study, two types of PLS were used. First, a mean-centered PLS approach was used to test for differences between groups (ELGA, VLGA, and Term). Accordingly, the contrast reflects the contribution of a group, the z-scores reflect the reliability of a given feature (a frequency-specific relative power value in a source location) and the p-value indicates the confidence in rejecting the null hypothesis of no group differences represented by the contrast.

Second, a behavioral PLS approach where the features (frequency power at a location) were investigated regarding its correlation with continuous variables (neurocognitive, neonatal, volumetric or intensity variables). In this case, the contrasts reflect the contribution of each variable, the z-score the reliability of the feature correlating with the set of variables, and p-values indicate the confidence in rejecting the null where the set of variables are randomly correlated with the features. To increase statistical power, in all the behavioral PLS analyses all the subjects from all the groups with information on the variable of interest were included in the analysis.

To assess associations between neuroimaging measures and neurocognitive outcome at school age, 11 variables were included representing IQ, CBCL, BRIEF and BEERY subscales; in the intensity correlation, the included variables were thalamic, cortical gray matter and cortical white matter mean intensities; for the volumetric PLS, the thalamic volume (TV) and cortical gray matter volume (CGV) ratio was obtained by dividing the former by the latter, to obtain the TV-CGV ratio. This was done to investigate associations between MEG power and TV not explained by CGV, as we hypothesized that decreased thalamic volume with increased cortical mass might better capture power changes across frequencies due to a reduction of thalamic cortical regulation relative to the amount of cortical neural mass.

Given that the TV-CGV ratio correlation with power does not tease out the sole contribution of TV or CGV, the TV and CGV were correlated with the so called ‘brain scores’ from the mean-centered PLS analysis. The brain scores are the dot product of the subjects by features matrix with the left-SV, and it expresses the covariance between the subjects’ features and SV. Essentially, the PLS ‘brain score’ for each subject indicates the degree to which that subject contributed to the pattern observed in the group contrast. A significant correlation between a volume variable and the brain scores would provide evidence of their contribution to the mean-centered PLS group differences.

The second technique used was linear regression. Given that neonatal variables were not significantly correlated with power in the preterm group, a further analysis was conducted using linear regression to test if GA, sex, pain, morphine, number of days in ventilation, and illness (SNAP) were associated with TV-CGV ratio or with thalamic intensity, which in turn were associated with power and explained the association with negative neurocognitive outcomes and group differences, respectively.

## Results

### Analysis of group differences in resting neurophysiological activity

When testing for group differences in relative power between ELGA, VLGA and Term, a statistically significant group difference was found with a p-value < 0.05 and a data-driven contrast reflecting ELGA vs. VLGA and Term. As illustrated in Fig. 1, the strongest positive z-scores (increased power for ELGA) were found for frontoparietal delta, frontal theta, and occipitotemporal gamma, while strongest negative z-scores (decreased power for ELGA) were located in posterior areas in the alpha band and in frontal areas in the beta band. Overall, higher delta, theta and gamma and lower alpha and beta were associated with power in ELGA compared with VLGA and full-term children.

**Figure 1.**
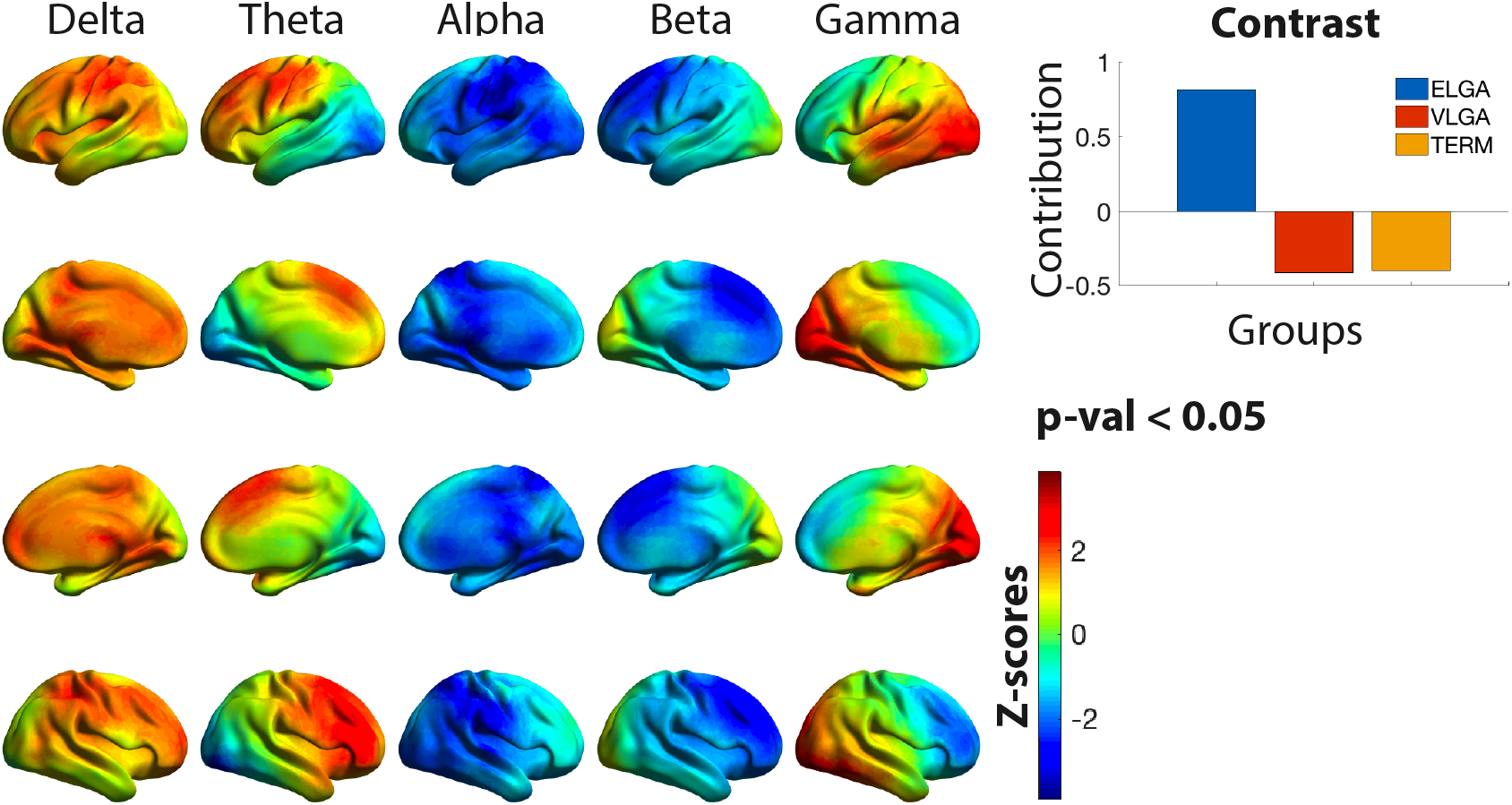
Group differences in power across frequency bands. Plotted on the cortical surfaces are z-scores indicating the reliability of expressing the group contrast plotted on the top right. The contrast indicates ELGA>VLGA and term, thus red z-scores reflects higher power in the ELGA group. Statistical analysis was performed on an 8 mm spaced grid, for visualization purposes, the z-scores where interpolated onto cortical surfaces.

### Power associations with neurocognitive measures

We tested for associations between power and a set of 11 neurocognitive measures and rendered a p-value < 0.05 and a data-driven contrast reflecting negative outcomes (CBCL, BRIEF) vs. positive outcomes (IQ, BERRY). The strongest z-scores, indicating more negative outcomes, reflected higher delta in general and theta in occipitotemporal areas and lower alpha, beta and gamma (Fig.2). In short, poorer outcome was associated with increased power at slower frequencies and decreased power at faster frequencies.

**Figure 2.**
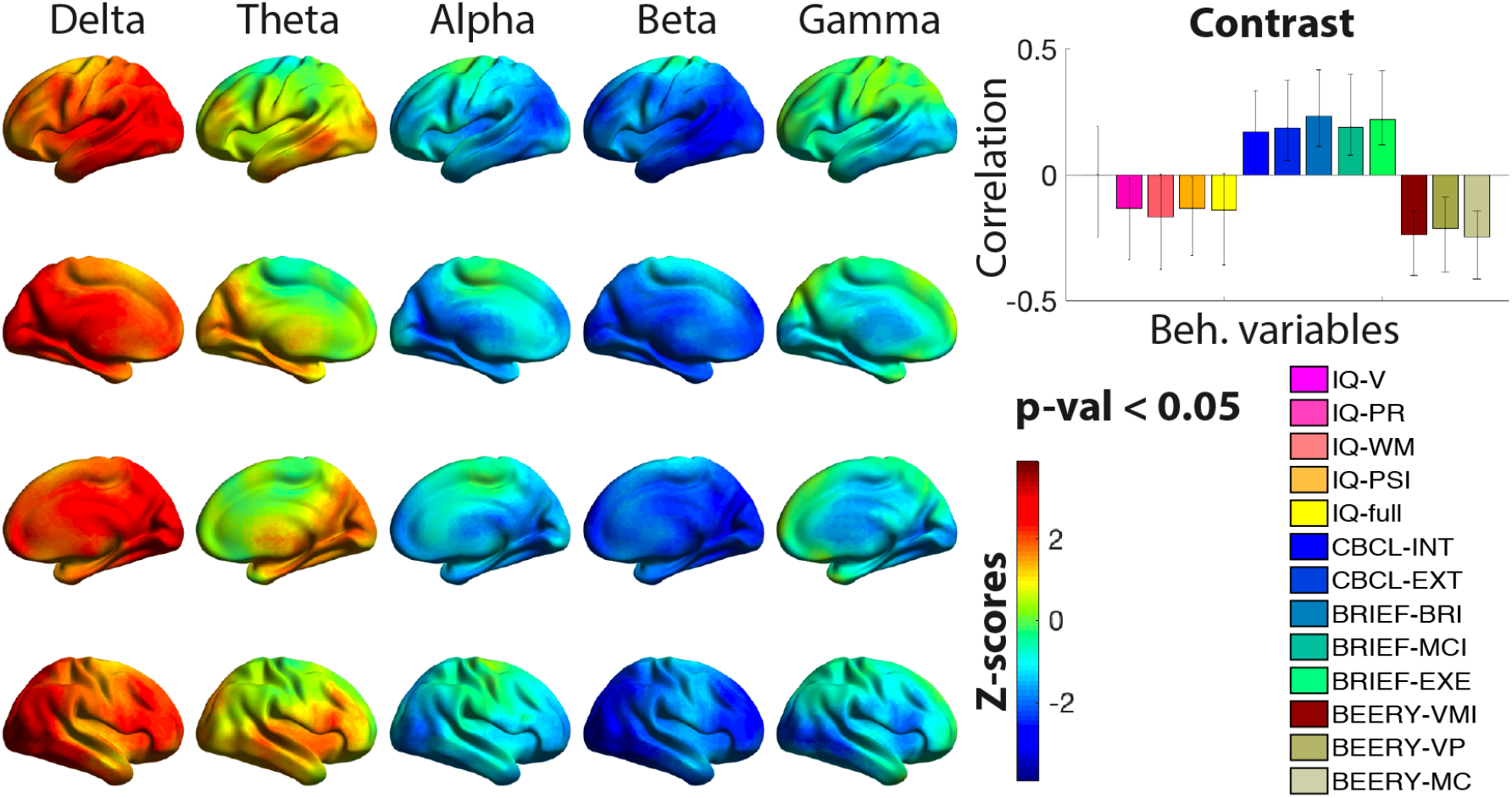
Correlation between power and neurocognitive measures. Plotted on the cortical surfaces are z-scores indicating the reliability of expressing the correlation between the set of neurocognitive variables (explained in Methods section) and power. The bar graph contrast plotted on the top right indicates the correlation values for each of the variables and the whiskers indicate the 95% confidence interval. The contrast indicates negative > positive outcomes, thus the higher the power in red z-scores, the higher the negative outcomes, i.e. lower IQ, higher difficulties in executive functions and lower visual motor integration. Statistical analysis was performed on an 8 mm spaced grid, for visualization purposes, the z-scores where interpolated onto cortical surfaces.

### Power associations with intensity measures

A significant association was found with a p-value < 0.05 between neurophysiological brain power and mean intensity variables. The data-driven contrast reflected higher cortical gray matter intensity (CGI) and cortical white matter intensity (CWI) vs. lower thalamic intensity (TI). Lower TI and higher CGI and CWI was associated with higher power in frontoparietal areas in the delta band, frontal theta and occipital gamma, and lower alpha and beta. The z-scores distribution resembled highly the PLS group differences, we correlated both z-scores and obtained a correlation value of 0.7.

### Associations between MEG power and volumetric measures

For PLS purposes, it would be equivalent to correlate power with TV and CGV as two separate variables, however, taking the TV-CGV ratio as a single variable allowed to further explore the relation with power and neonatal variables. PLS significantly correlated power with the TV-CGV ratio with a p-value < 0.01. Decreased TV relative to the cortical volume of gray matter was associated with higher delta in general and theta in occipitotemporal areas, and lower alpha, beta and gamma (Fig.4). The z-score distribution was very similar to the PLS association with neurocognitive measures, when correlating both z-scores we obtained a correlation value r = 0.9. *A posteriori*, to further assess if the results were driven by either TV or CGV, a separate PLS was conducted, and only the TV showed a significant trend with a p-val <.1. To tease out if the ratio reflects the contribution of both variables or if it is driven by the TV while accounting for CGV, pearson correlation with either TV or CGV and the brain scores from the mean-centered PLS (group differences in power) was calculated. The correlation value for TV and brain scores was r=.4 and a p-val < 0.01, while CGV correlation was not significant, suggesting the TV as the main contributor.

**Figure 3.**
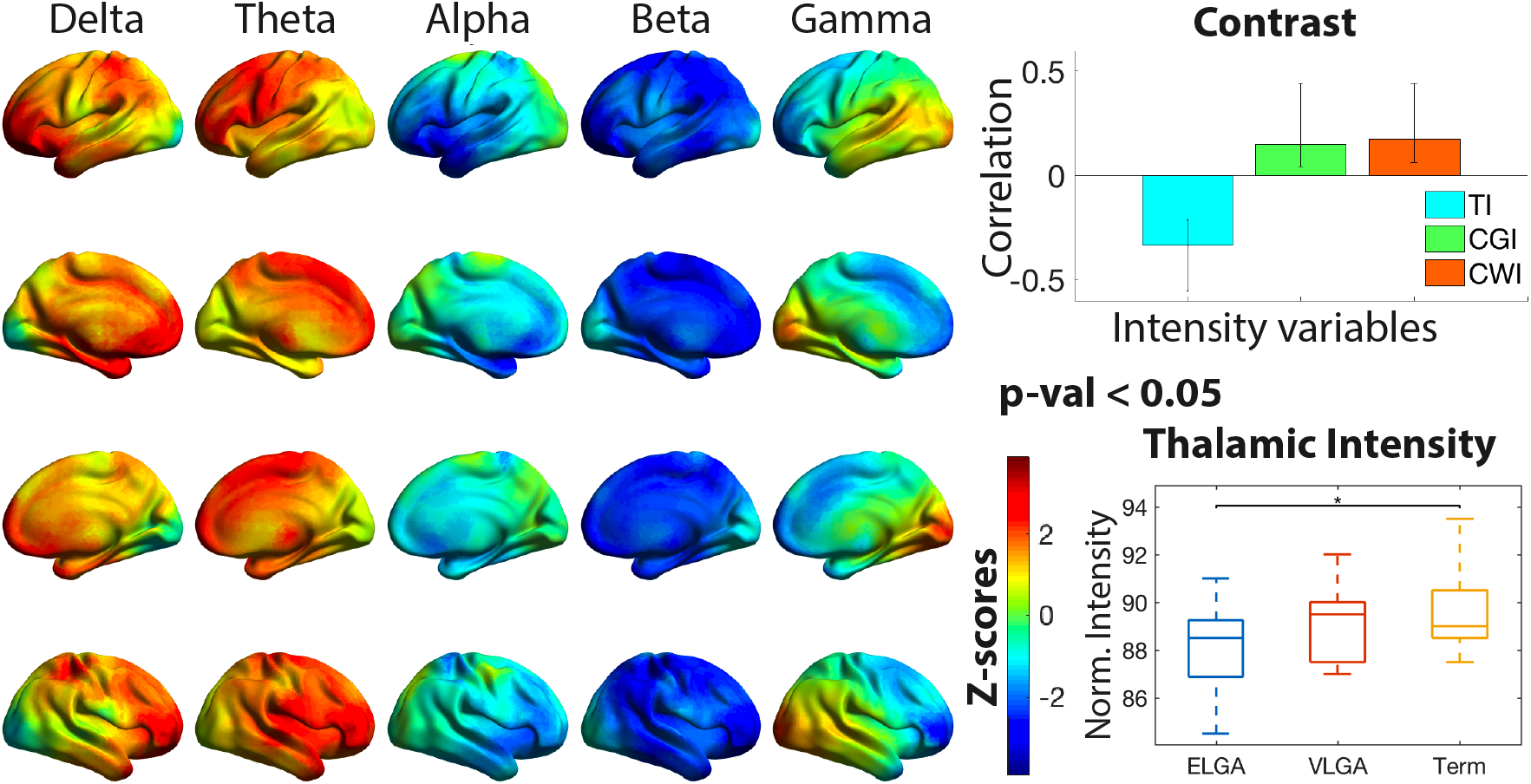
Correlation between power and normalized mean intensity measures. Plotted in the cortical surfaces are z-scores indicating the reliability of expressing the correlation between the set of intensity variables (mean intensity in the thalamus (TI), cortical gray matter (CGI) and cortical white matter (CWI), explained in Methods section) and power. The bar chart contrast, plotted on the top right, indicates the correlation values for each of the variables, and the whiskers indicate the 95% confidence interval. The data-driven contrast indicates lower TI vs. higher CGI and CWI, thus the higher the power in red z-scores, the smaller the TI and higher the CGI and CWI. Statistical analysis was performed on an 8 mm spaced grid, for visualization purposes, the z-scores where interpolated onto cortical surfaces. On the bottom right, the TI values are plotted with a box plot for each group, horizontal lines indicate significance between groups, * = p-val <0.05.

**Figure 4.**
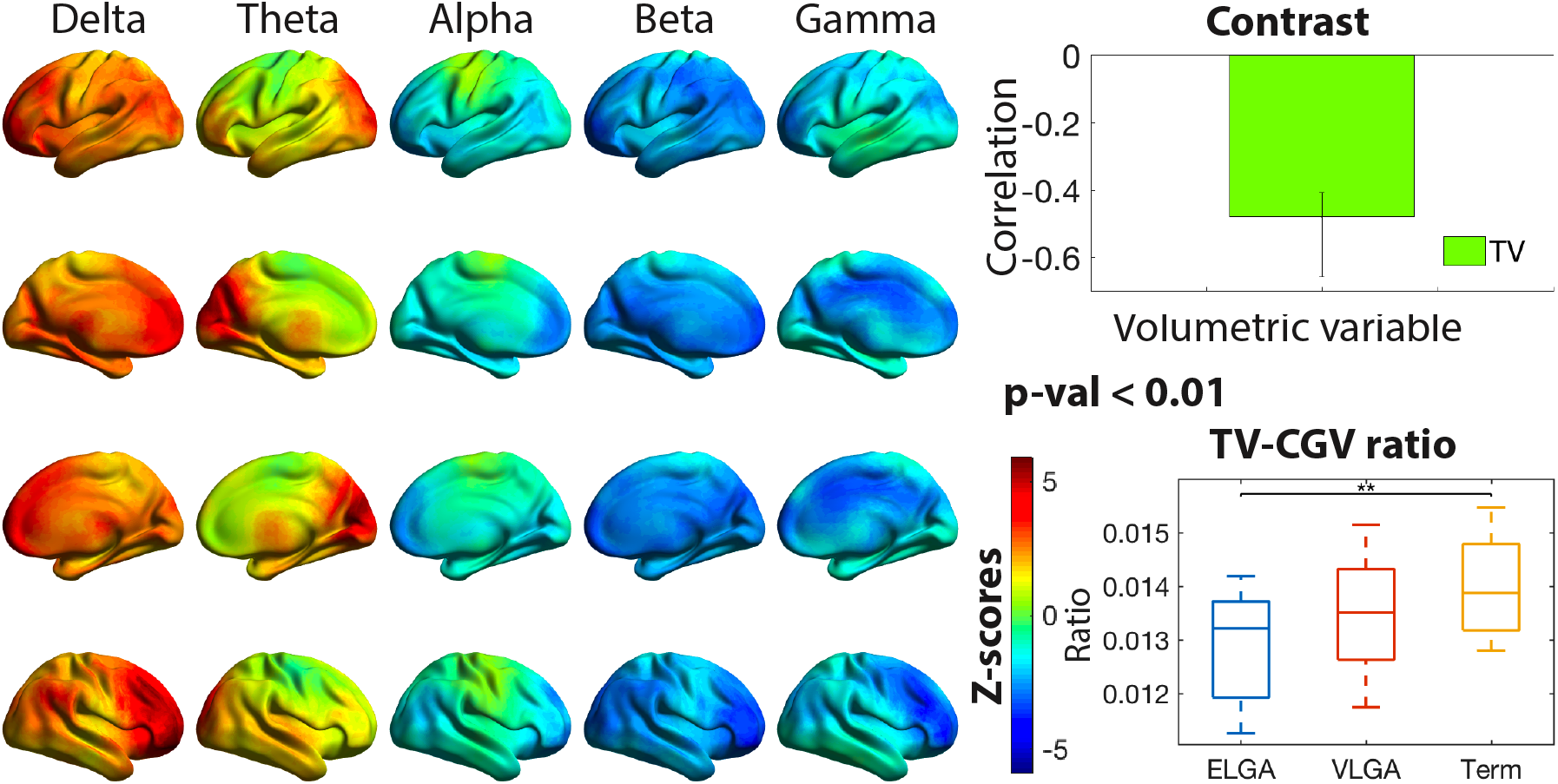
Correlation between power and the thalamic and cortical gray volumes ratio. Plotted in the cortical surfaces are z-scores indicating the reliability of expressing the correlation between the thalamic and cortical gray volumes (TV-CGV) ratio and power. The bar chart contrast, plotted on the top right, indicates the correlation value for the TV-CGV measure, and the whiskers indicate the 95% confidence interval. The contrast indicates > TV-CGV, thus the higher TV-CGV the higher the power in red z-scores areas. Statistical analysis was performed on an 8 mm spaced grid, for visualization purposes, the z-scores where interpolated onto cortical surfaces. On the bottom right, the TV-CGV values are plotted with a box plot for each group, horizontal lines indicate significance between groups, ** = p <0.01.

We explored the association between the average power from the 2.5% locations with the strongest z-scores in each frequency band. For each frequency, a scatterplot and the line of best fit for each group was plotted and correlated the ratio with the average power for all the participants and within group (Fig. 5). The scatterplots and correlations indicate that the association of TV-CGV ratio and power is present across all frequencies and groups with the same effect direction, although, by combining all the groups higher statistical power is obtained.

**Figure 5.**
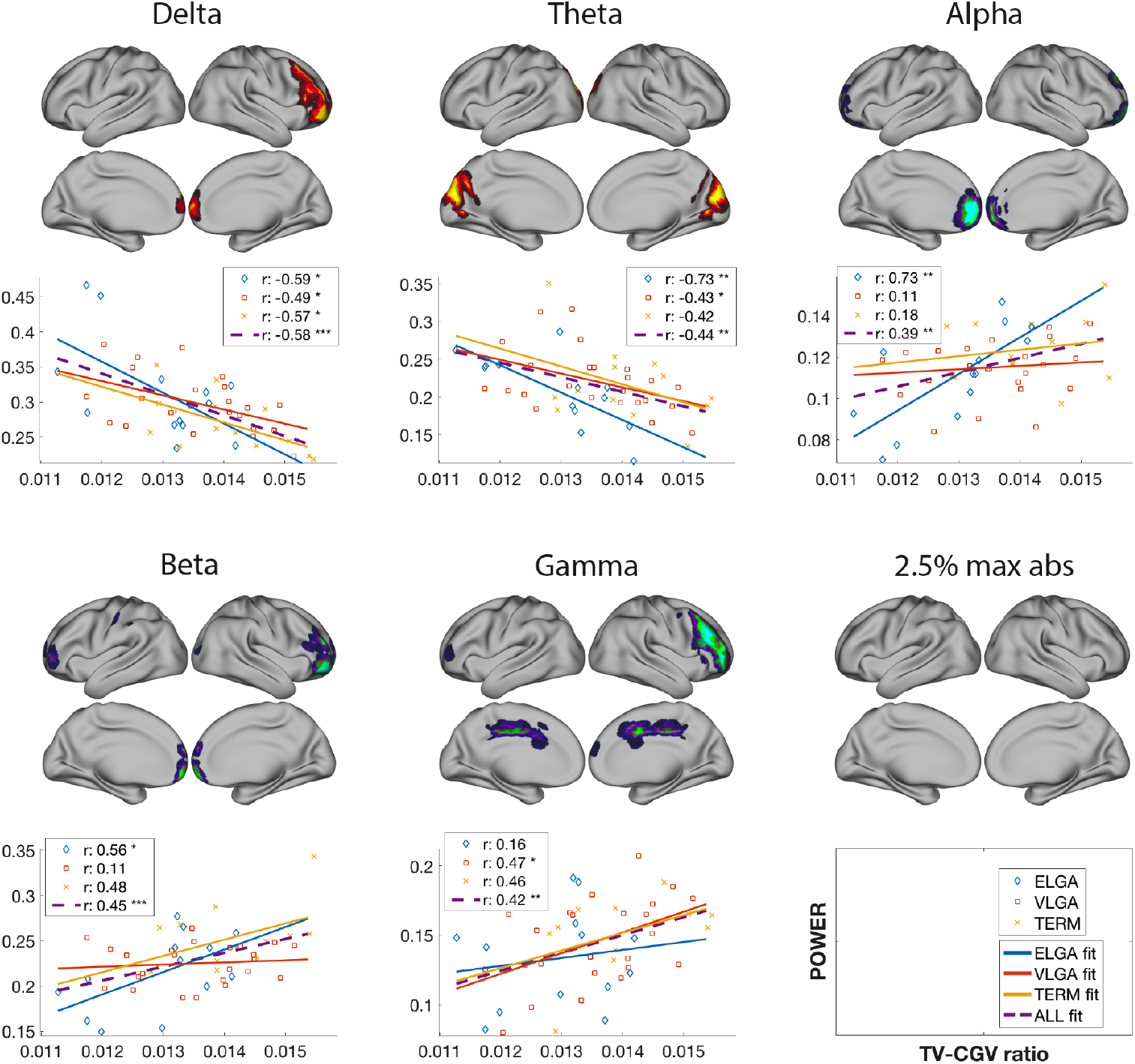
Correlation between average power from locations with the highest magnitude z-scores and the TV-CGV. Plotted in the cortical surfaces are the areas with the biggest absolute z-scores thresholded at 2.5%. For delta and theta frequency bands, the z-scores were negatively skewed, whereas in the other bands were positively skewed with predominantly more negative z-scores. The 2.5 tail from the skewed side (plotted on the brain surfaces) at each frequency were selected and their power averaged. The averaged power was correlated with the TV-CGV ratio, for each group and in total, and plotted in a scatterplot. The lines in the scatterplot represent the best linear fit, and in the box legend the correlation and significance is presented, with *= p<.05, **= p<.01, ***= p<.001. In all the groups and frequencies, the direction of the association (positive or negative) is common across groups, power in ELGA tends to correlate more, but overall, the association with all the participants is highest.

### Neonatal variables associations with volume and intensity

Two linear regressions were calculated to predict either the TV-CGV ratio or the TI based on neonatal variables. The model for the TV-CGV was not significant, whereas the TI significance was p<0.05, F(7,29)=2.65, , with R^2^.37. Beta and p-values for the predictor variables are reported in Table 2.

**Table 2.** Linear regression for thalamic mean normalized intensity. Predictor variables are gestational Age (GA), sex, GA*sex interaction, infection, illness severity (SNAP), cumulative morphine dose, number of skin breaking procedures (pain). P-values in red indicate statistical significance.

|  | Beta | Sig. |
| --- | --- | --- |
| GA | -0.104 | 0.616 |
| Sex | -14.14 | 0.030 |
| Infection | -1.084 | 0.063 |
| SNAP | -0.009 | 0.729 |
| Morph | -2.948 | 0.008 |
| Pain | -0.064 | 0.954 |
| Sex*GA | 0.487 | 0.027 |

## Discussion

This study examined relationships between frequency-specific power, structural alterations associated with the thalamus and neurocognitive difficulties at school age that characterize children born very preterm. We found that relative power at rest in the extremely low gestational age (ELGA) group differed the most compared to the very low gestational age (VLGA) group and term control group, as similarly reported with absolute power [Kozhemiako et al., 2018], and likely related to prior observation of reduced alpha connectivity during task performance [Doesburg et al., 2010]. Neurocognitive difficulties were associated with increased frontal and occipital delta, and occipital theta, as well as decreased power mostly involving frontal and occipital beta activity. Thalamic, cortical gray matter and (sub)cortical white matter intensities were associated with power and were very similar, in direction and spatially, to the group differences in power, differentiating ELGA from the rest, with a correlation between z-scores of r=.7. Similarly, the association between MEG power and the ratio thalamic/cortical volume revealed a spatial pattern very similar to that observed in the association between MEG power and neurocognitive difficulties. These similarities suggest that the thalamic intensity, accounting for cortical gray and white matter intensity, is a prominent structural correlate of atypical power in the ELGA, and the thalamic volume, normalized by the total neural cortical mass, is the structural correlate of neurocognitive difficulties. Overall, this study provides novel evidence of the relationship between atypical power at rest in preterms and structural thalamic alterations and negative outcomes at school age.

The central hypothesis of this study was based on reports indicating thalamic volume reduction and thalamocortical and corticothalamic pathways alterations in children born very preterm [McQuillen and Ferriero, 2006; Smyser et al., 2010], on studies suggesting the role of the thalamus as a regulator of cortical through low frequencies [FitzGerald et al., 2013; Malekmohammadi et al., 2015; Ribary, 2005], and on studies demonstrating the importance of the thalamus for higher-order cognitive processes [Mitchell, 2015; Saalmann, 2014; Sweeney-Reed et al., 2015]. Based on thes previous literature, we hypothesized that decreased thalamic volume might lead to abnormal distribution of power at different frequency bands which would be associated with negative neurocognitive outcomes. Indeed, we found that atypical power in the ELGA group marked by a relative power increase of lower frequencies (delta and theta) and decreased higher frequencies (alpha, beta, and partially gamma) was also associated, in general, with more negative outcomes. These results are consistent with the phenomena of ‘alpha slowing’, which is characterized by decreased alpha power and increased theta and/or delta power [Garcés et al., 2013; Hughes and Crunelli, 2005; Llinás et al., 1999]. This slowing has been associated with a decrease of inhibitory interneurons in the thalamus or a shift of inhibitory drive on the thalamus [Bhattacharya et al., 2011; Ribary et al., 2019], and based on our results, this appears to be structurally reflected with decreased gray matter intensity in the thalamus. This power imbalance seems to be already present in the neonatal period in preterm infants with a relatively higher contribution of low frequencies [Suppiej et al., 2017]. Similarly, Doesburg et al. 2011, using a subset of the participants analyzed in the present study, found on sensor-space slowing of alpha peak especially in the ELGA children, during a visual-spatial memory task at age 7-8 years. In the present study, we show a source-space spatially defined characterization of atypical power, most distinctly evident in children born extremely preterm, and the association with structural thalamic deficits and neurocognitive difficulties.

In the analysis of associations between power and thalamic volume (TV) or intensity (TI), *a priori*, the cortical counterpart was included, i.e. cortical gray matter volume (CGV) or intensity (CGI), to account for power independent from thalamic alterations. Although it has been found decreased cortical thickness in preterms at school age [Lax et al., 2013; Ranger et al., 2013], very preterm birth is associated with neurodevelopmental problems reflected in slower neuronal optimization archived by pruning unnecessary synapses and strengthening necessary ones with myelinization [Shaw et al., 2008]. Longitudinal preterm studies have found that between 7 to 12 years of age there is decreased thinning of cortical gray matter volume indicating delayed cortical maturation driven by cortical thinning [Ment et al., 2009; Mürner-Lavanchy et al., 2014]. In our study, while CGV was significantly reduced in the ELGA, we found that the ratio TV-CGV was also significantly decreased, indicating a relatively bigger CGV. In relation to the thalamus, we hypothesize that reduced thalamic regulation, either by a smaller number of thalamic neurons or pathways, might lead to cortical dysfacilitation were more cortical synapses are needed to compensate for the reduced role of the thalamus in integrating cortical processing. This hypothesis would also account for the increase of delta and theta seen in the ELGA, as more low-frequency integrative processing in the cortex would be needed, and, assuming cortical integration to be less effective than the thalamic, it would also explain the correlation between neurocognitive difficulties and higher low frequency power.

We expected to find a direct association between power and neonatal procedures, especially, pain (skin breaking procedures). However, at most, we found an indirect link with power by using neonatal variables as predictors of TV-CGV ratio or thalamic intensity (TI). While TV-CGV was not significantly associated, TI reduction was predicted by morphine instead of pain, with a more pronounced effect on females. Gray matter intensity mostly reflects the amount of myelinization or the amount of axonal mass. While subcortical white matter intensity reflects axons from the thalamus and elsewhere, thalamic gray matter intensity reflects structural connections involving directly the thalamus. Preterm alterations in myelinization have been previously reported [Lubsen et al., 2011; Nossin-Manor et al., 2013], including in the thalamus [Eaton-Rosen et al., 2015], with early exposure to opioid affecting myelination [Sanchez et al., 2008], nonetheless, the prediction of morphine for TI might also reflect its apoptotic effects in the thalamic neurons and microglia [Hu et al., 2002; McPherson and Grunau, 2014]. In this study, we provide evidence that higher cumulative dose of morphine is a predictor of TI which in turn is associated with atypical power in ELGA. This is consistent with previous findings of adverse effects of high dosing of morphine in relation to hippocampal [Duerden et al., 2016] and cerebellar [Zwicker et al., 2016] growth on MRI in the neonatal period in a different cohort, and cerebellar growth in the current cohort at school-age [Ranger et al., 2015].

In conclusion, our findings present evidence linking atypical relative power at rest in children born very preterm, especially ELGA, at school age with neurocognitive difficulties, structural deficits related with the thalamus, and neonatal pain and morphine exposure. During the neonatal period surrounding very preterm birth, the thalamocortical system is going through critical developmental windows very susceptible to disruptions that might lead to long-lasting alterations, at least until school age. The present work raises the issue of the effects of preterm birth, as well as the importance of optimizing neonatal procedures and the possible integration of early cognitive interventional training programs in early childhood.

## ACKNOWLEDGEMENTS

We would like to thank Dr. Ken Poskitt, pediatric neuroradiologist, for his invaluable assistance in obtaining the MRI scans. We would like to thank Dr. Ivan Cepeda and Gisela Gosse for coordinating the study, Katia Jitlina, Amanda Degenhardt, Dr. Teresa Cheung and Julie Unterman for their help in data collection. This study was supported by Grant R01 HD039783 from the Eunice Kennedy Shriver Institute of Child Health and Human Development (REG), the Canadian Institutes of Health Research Grants MOP42469 (REG) and MOP-136935 (SMD).

## CONFLICTS OF INTEREST

The authors declare no conflicts of interest.

